# Molecular Dynamics Simulations Studies On The Effects Of Mutations On The Binding Affinities Between SARS-CoV-2 Spike RBD And Human ACE2

**DOI:** 10.1101/2022.02.11.480063

**Authors:** Smriti Arora, Jeevan Patra

## Abstract

The SARS-CoV-2 viruses had made a great impact on humankind and the world economy. Phylogenetic analysis revealed the newly identified B.1.617.1 and B.1.617.2 lineages possessed with few key mutations predominantly circulating. The signature mutations possessed by these lineages are situated in the RBD motif of S protein. Reports revealed variants L452R, T478K, and E484Q harbours in enhancement with hACE2 binding while P681R situated in furin cleavage site resulting in better transmissibility. To gain a deeper understanding of the impact of these variants (L452R, T478K and E484Q) binding with hACE2, structural dynamics at the interface between S-RBD protein and hACE2 were studied. We performed our dynamics studies with both single mutant complex (L452R, T478K and E484Q) and in the combination of triple mutants (L452R + T478K + E484Q) at 100ns in contrast with the wild type. Interfacial docking interactions and Molecular Mechanics approach exhibited that the spike mutants −L452R, T478K and E484Q harbour with higher binding affinity on hACE2 in contrast with its native spike protein. The presence of interfacial residue, intermolecular contacts such as hydrogen bonding, salt bridge and non-hydrogen bonded interactions might be the reason for its higher binding affinity. Hence the findings from our study unravelled plausible mechanism for the increase in affinities of mutants to hACE2 thus leading to higher transmissibility and infection of emerging variants. Further, the conformational alterations in the course of dynamics at the RBD motif led to enhancement of hACE2 binding and immune escape. These results suggest that the structural changes introduced by these variants enhance the binding affinities of the S protein with the hACE2 that could form the basis to further aid in designing therapeutics that could inhibit at the interface of S protein and hACE2 receptor.

## BACKGROUND

The COVID-19 caused by the Severe Acute Respiratory Syndrome Coronavirus 2 (SARS-CoV-2) have made a calamitous impact on worldwide. Since its emergence from December 2019, total number of the cases hiked to more than 220 million and of overall 4 million causalities. The influence of mutations caused tremendous impact towards the health in challenging 2020-2022 year. The accelerated development of therapeutics (drugs and vaccines) with impressive efficacy and potency by the researchers fulfilled unmet needs to the humankind (Yan et al., 2021). Despite all the efforts and hype, the incarnation of new viral variant of concerns from the lineage B.1.617 exhibited to have high transmissibility to escape the immune system from the hosts. The cellular mechanism of Severe Acute Respiratory Syndrome coronavirus 2 (SARS-CoV-2) surface spike proteins engulf into host human cells and interacting to the angiotensin-converting enzyme-2 (ACE2) with the viral receptor binding domain (RBD) (Shang, Ye, et al., 2020). The available crystallographic structure of spike subunits of SARS-CoV-2 or its RBD complex with host ACE2 indicated that the presence of the core in RBD and Receptor Binding Motif (RBM) which interacts with the ACE2 (Shang, Ye, et al., 2020). From the thorough literature survey, the mutagenesis in the RBM responsible for the infectivity, transmissions, aetiology and immune escape (Wan et al.). The aetiology of SARS-CoV-2 remains unclear. The single positive stranded RNA virus comprises of four major structural proteins, such as spike, nucleocapsid, envelope and membrane protein. The spike protein is a large oligomer transmembrane that mediates viral entries into the host cells. It is composed of two subunits S1 and S2 for receptor binding and membrane fusion respectively (Walls et al., 2020; Wrapp et al., 2020). The spike protein of the virus exploits hACE2 protein for invading into the host cells. Hence, S protein represented as the infectivity and transmissibility into the host (Hulswit et al., 2016; Letko et al., 2020). Studies revealed that the S1 subunit (RBD) mediates through ACE2 during the virus interactions. This interaction occurs in between S1-S2 junction by furin cleavage enzyme expressed by the host cells, which is further transmitted, to S2 subunit for efficient infection (Hoffmann et al., 2020; Shang, Wan, et al., 2020) [**Figure 1**].

**Figure 1:**
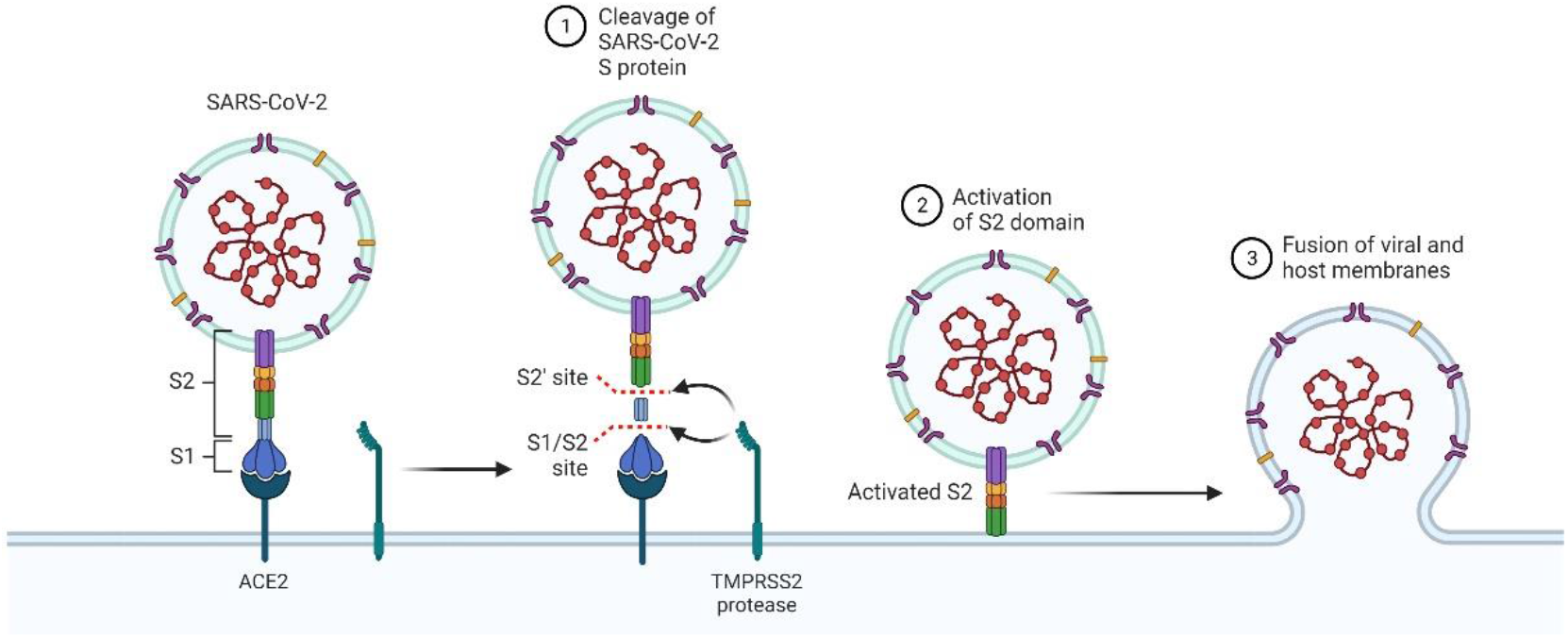
Mechanism of SARS-CoV-2 spike protein invading to host cells

As the global pandemic health burden emerges, the completely genomic sequence been considered to analyse and understand the transmissibility and its evolution. Reports revealed that the D614G mutant in the spike proteins found to have significantly higher human host infectivity and transmission efficiencies of the SARS-CoV-2 from host-to-host (Korber et al., 2020; Volz et al., 2021; L. Zhang et al., 2020). Recently, three new variants of concerns (VOCs) emerged such as B.1.1.7-α, B.1.351-β and P.1-γ in the RBD region such as N501Y, E484K and K417N/T (Leung et al., 2021; Rambaut et al., 2020; Volz et al., 2021). Upon the influence of several mutations, viruses continuously adapt new forms where the mechanisms responsible for the emergence of new variants. The viruses consist with the RNA found to have enormous stakes of mutation levels in comparison with the DNA viruses (Duffy, 2018; Lauring & Andino, 2010). The presence of the amino acids on the virus surface significantly alters its virus functions, results in transmissibility and mortality (Diehl et al., 2016; Herfst et al., 2012; Tsetsarkin et al., 2007). In SARS-CoV-2 the gene encoding with the spike protein, found to have several mutations those were reported earlier such as K417N, E484K, and N501Y (Dawood, 2020; Korber et al., 2020). The borne of this amid crisis in South Africa and United Kingdom have faced enormously based on the new variants as named 501Y. V2B.1.351 and 501.V2 for South Africa and UK respectively. Earlier reports revealed that these new variants harbours in multiple regions of the spike protein where the point site mutation of specific amino acids replacements are K417N, E484K and N501Y for the South African strains and N501Y for the UK strains at the key region of the RBD domain (Andrew Rambaut, 2020; Tegally et al., 2020).

Upon the second wave from March 2021, there were a tremendous sharp rise in COVID-19 cases and deaths in India. From the phylogenetic analysis the viral variant in India belonging to the newly identified B.1.617 lineage. This lineage has received great attention due to its increased infectivity, high virulence capacity, and potential immune escape. This lineage has three sub-categories such as B.1.617.1, B.1.617.2 and B.1.617.3 harbouring on the spike protein in the viruses. The mutation variations bearing with this lineage are D111D (synonymous) and G142D, L452R, E484Q, D614G and P681R (non-synonymous). The combination of these mutations provides virus survival and the ability to spread rapidly. The structural insights from the available crystallographic Spike-RBD-ACE2 complex protein, both these E484Q and L452R variants harboured in the Receptor Binding Motif (RBM) of the S protein.

Gaining deeper insights on the interactions between the viral S protein and hACE2 is essential for developing therapeutics as well as finding the efficacy of existing vaccinations against SARS-CoV-2 viruses. In our research, we gave an attempt to gain novel insights on the structural sites in between ACE-2 and RBD protein interactions using hyphenated silico techniques. We predominantly focused on the mutations located on spike RBD domain with the emphasis on the potential impact on three variants (L452R, T478K and E484Q) that are directly located at the hACE2 interface. In addition, we extended our investigations on the protein structural stability, binding affinity and intermolecular interactions in between the S protein and hACE2 (L452R, T478K and E484Q, independently) and in triple combinations (L452R + T478K + E484Q).

## COMPUTATIONAL METHODS

### Protein Preparations and *in-silico* Mutagenesis

The x-ray crystallography of the Spike-RBD-ACE2 complex (PDB 6M0J) with resolution of 2.45 Å was retrieved from protein data bank (Lan et al., 2020). The protein preparation was performed with these mutant complexes which involves bond order assignment, creating disulfide bonds, adjusting ionization states, removing water molecules and hetero atoms, metals and co-factors, correction of group orientation and termini capping, adding missing atoms and side chains ads partial charges assignment. The hydrogen bonds were optimized using PROPKA at neutral pH. From the wild type, three mutations such as L452R, T478K and E484Q by performing key replacement of residues using mutagenesis module of PyMOL v2.5, Schrodinger LLC (DeLano, 2002). The rotamers were considered based on highest frequency of occurrence. All the stability and flexibility from the obtained trajectories from were examined using CABS-Flex. CABS-Flex – a standalone fast modelling package was considered to understand the flexibility of both the Spike RBD domain and ACE2-Spike complex (Badaczewska-Dawid et al., 2020; Kurcinski et al., 2019; Kuriata et al., 2018)

### Molecular Dynamics Simulations and MM-GBSA calculations

The molecular dynamics simulations were performed up to 100ns using Desmond package (Bowers et al., 2006; Chow et al., 2008). Both the wild and mutant complexes were placed in orthorhombic box at absolute size and solvated using Single Partial Charge (SPC) using Desmond system builder. The simulation system was neutralized with counter ions and salt concentration of 0.15 M NaCl. All complex were described with the OPLS force field that were assigned to each simulation run for 100ns. The dynamics simulation was maintained with the isotropic Martyna-Tobias-Klein barostat and Noose-Hoover thermostat at 1 atm and 300K pressure and temperature respectively (Martyna et al., 1992; Martyna et al., 1994). The smooth Particle Mesh Ewald method used for evaluating long range coulombic interactions with a short-range cut-off at 9.0 Å (Essmann et al., 1995). The binding free energies of all proteinprotein complexes with individual residue free energy contributions were computed and estimated with help of Molecular Mechanics Generalised Born Surface Area (MM-GBSA) using HawkDock (Weng et al., 2019). All the pictorial representation was drawn and retrieved using Visual Molecular Dynamics – 1.9.3 (Humphrey et al., 1996).

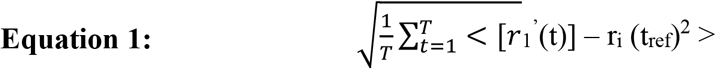

Where, T is the trajectory time over which the RMSF is calculated, t ref is the reference time, r i is the position of residue i; r’ is the position of atoms in residue i after superposition on the reference, and the angle brackets indicate that the average of the square distance is taken over the selection of atoms in the residue.

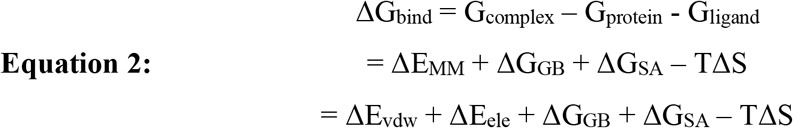

Where, ΔE_MM_ is the interaction energy with two phases ΔEvdw (van der waals energy) and ΔEele (electrostatic energy) (Genheden & Ryde, 2015). ΔG_GB_ and ΔG_SA_ are the de-solvation free energy (Onufriev et al., 2004). Normal mode analysis used for evaluating entropy contribution to the binding free energy.

### Protein-Protein Probing and Interfacial Analysis

The contacts in between hACE2 with three mutant complexes were studied with High Ambiguity Driven Protein-Protein DOCKing (HADDOCK) (Honorato et al., 2021; van Zundert et al., 2016). The binding sites were defined within 3Å at the interfacial region based on the interaction profiles of wild type (PDB 6M0J). All the conformations were ranked based upon the HADDOCK PPI score.

HADDOCK Score = 0.2 × Electrostatic energy + 1.0 × Van der Waals energy + 1.0 × De-solvation energy + 0.1 × Restraint’s violation energy

The binding potential energies and protein-protein interface interactions was estimated using mCSM-PPI2 (Rodrigues et al., 2019). All the interactive analysis were performed using UCSF Chimera and PyMol visualizer (DeLano, 2002; Pettersen et al., 2004).

## RESULTS AND DISCUSSIONS

SARS-CoV-2 viruses, member of coronaviridae family, betacoronavirus genus and sarbecovirus subgenus with having 29.9-kb linear single stranded positive sense RNA genome (Lu et al., 2020; M. Y. Wang et al., 2020). The S glycoprotein composed of S1 and S2 subunits found on the surface which aids in the cellular hACE2 host cells affinity and membrane viral fusion respectively (Satarker & Nampoothiri, 2020; H. Zhang et al., 2020). The COVID-19 has caused drastic impacts on people’s daily lives, and world economy (Li et al., 2021). Structural insights proven that S1 subunit recognized as hotspot for mutations, virulence transmissibility, and immunological evasions in the hosts (Shang, Ye, et al., 2020; Yi et al., 2020). SARS-CoV-2 viruses evolved with numerous co-circulating variants of concerns since December 2019. From these variants, B.1.1.7 (alpha), B.1.351 (beta), P.1 (gamma) and B.1.617.1 and B.1.617.2 (kappa and delta) (Aleem et al., 2022). Earlier studies revealed that the binding ability of the variants could enhance in virus replication and transmissibility (Korber et al., 2020; Thomson et al., 2021). In this study, we took an attempt to gain deeper understanding in binding interactions of wild and mutant type (L452R, T478K, and E484Q) to hACE2 cells. Further, we tried to elucidate its plausible binding affinities mechanisms using interface docking analysis and adaptive molecular dynamics simulations. From the in-depth analysis, the binding energies of mutants exhibited higher affinity to hACE2 in contrast with wild type. The increase in binding affinities is correlated with the interfacial interactions such as hydrogen bonding, salt bridges and hydrophobic contacts. Our molecular dynamics simulation results gave insights into geometric parametric molecular aspects how affinity enhanced in spike mutants to hACE2 receptor, which supports for designing therapeutics targeting at RBD region of S glycoprotein.

The key roles of the amino acids in proteins are crucial for the regular cellular activities. If any alterations of the amino acids led to enormous impact on the protein structure and bio-molecular ligand interactions. Few structural parameters govern for the mutagenesis, such as the key replacements of the residues, region of the structure (loop or C-α backbone, shallow cavity or solvent exposed), and active site (catalytic or conserved) or interface of two dimers or protein-ligand (Gyulkhandanyan et al., 2020; Ittisoponpisan et al., 2019; Kucukkal et al., 2015; López-López et al., 2020; Thusberg & Vihinen, 2009). The growth of the bioinformatics defined all the strength and weakness of the available computational tools to understand the structural insights for the study of the mutagenesis and its structural functions when we have a crystallographic structure (Singh et al., 2021). Although we have vast resources for the protein understandings, however it is still recommended to gain insights using other algorithm designed tools to have consensus knowledge about the key amino sites replacements (Gyulkhandanyan et al., 2020). Our investigational study based on the reported data and its influence on the hACE2 protein (Starr et al., 2020). To initiate our study, we undergone thorough study in the Protein Data Bank. The crystallographic structures with ACE2 and RBD bounded complex been published (Lan et al., 2020; Q. Wang et al., 2020; Yan et al., 2020). The literature data also reflected that the polymorphism in the ACE2 gene curtails the binding affinity and low vulnerable towards the infection (Alifano et al., 2020; Calcagnile et al., 2021).

The proteins flexibility trajectories obtained from all three complexes – single mutation (L452R, T478K, and E484Q) and in combination triple mutation (L452R + T478K + E484Q) were analysed. The RMSF plots indicated that these systems marked its equilibrium and all three protein complexes are stabilized in all simulations [**Figure 2**]. Further analysis on the individual residual fluctuation and protein flexibility at Cα backbone atoms in between the spike – hACE2 interface, RMSF trajectories were computed and obtained. To gain deeper understanding and the necessity of the backbone loops while binding to hACE2, we had in-depth structural insights on the binding interface reported by Lan and his co-worker (Lan et al., 2020). From their study, interface comprises of four loop regions. Loop 1 (residues 438 – 450); loop 2 (residues 455 – 470); loop 3 (residues 471 – 491) and loop 4 (residues 495 – 508). Previously, study revealed that the loop 3 and 4 regions are more flexible domains at the RBD (Williams et al., 2021). Upon the closer observation, the residue flexibility trajectories also reflected to have higher flexibility at the region of 470 – 490.

**Figure 2:**
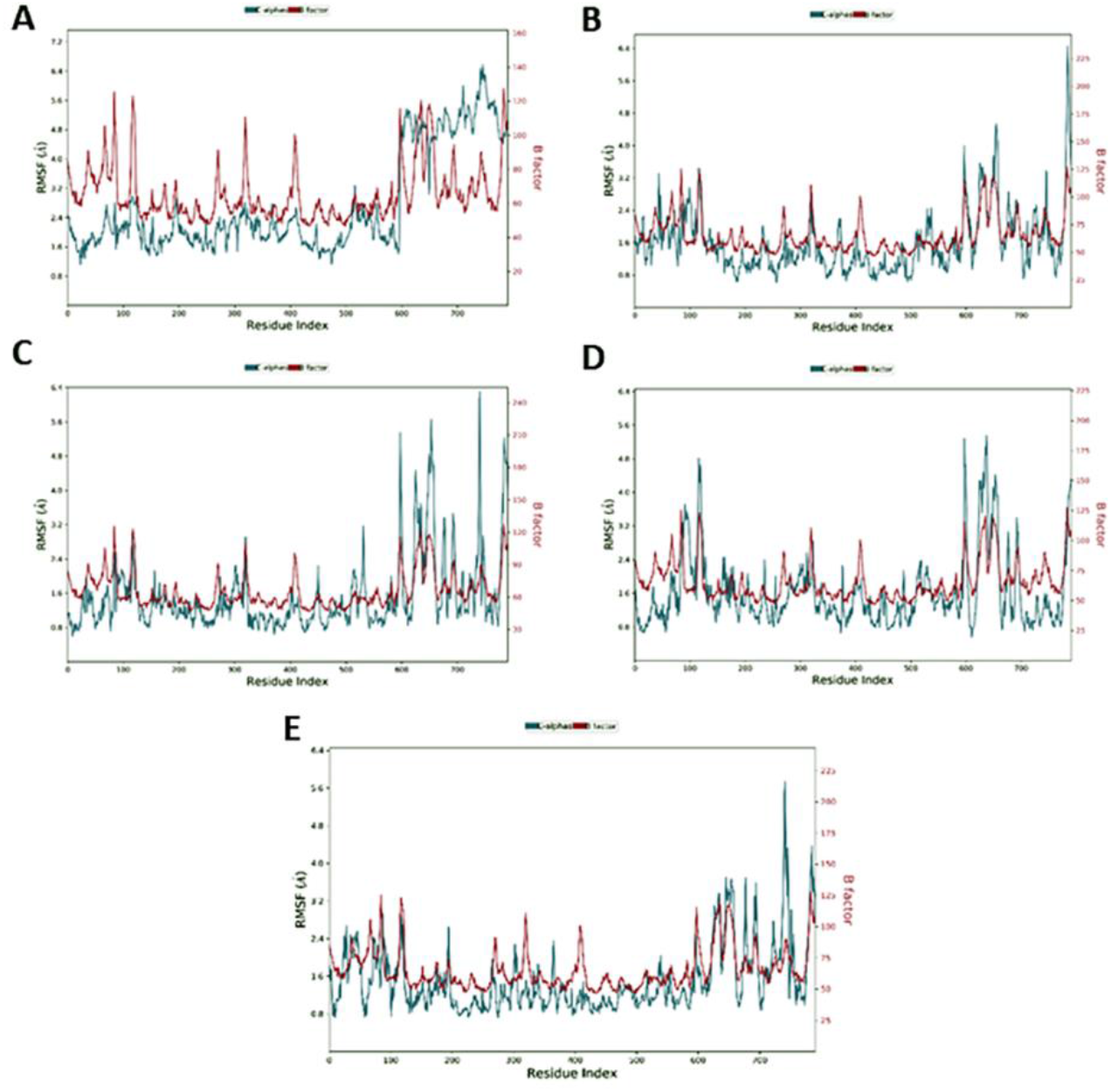
Root mean square fluctuation (RMSF) of Cα atoms (dark green) and B factor (red) of mutant complexes obtained at 100 ns simulations of SARS-CoV-2 spike protein bound to hACE2. (A) RMSF of S-RBD and hACE2 protein in the L452R complex; (B) RMSF of S-RBD and hACE2 protein in the T478K complex; (C) RMSF of S-RBD and hACE2 protein in the E484Q complex; (D) RMSF of S-RBD and hACE2 protein in the triple combination (L452R + T478K + E484Q) complex; (E) RMSF of S-RBD and hACE2 protein in the wild type.

During our simulation studies, several intermolecular hydrogens bonding, hydrophobic and other contacts observed to have interactions, cleavage of contacts and reformation on newer ones. Previously, study displayed to have four interfacial contacts of S protein such as Lys417, Gln493, Tyr449 and Gln498 with hACE2 such as Asp30, Glu35, Asp38 and Lys353 (Ali & Vijayan, 2020). Further, these results were performed comparative study with of previously reported wildtype results (Ali & Vijayan, 2020). The contacts exhibited from the interfacial analysis of all the single mutant complexes formed about 2Å (Figure 3). In the complexes L452R, T478K and E484Q single hydrogen bonding formed with hACE2 Glu35 at 2.1 Å, single salt bridge with hACE2 Glu75 at 2Å and two hydrogen bonding with Glu75 at 1.9 Å respectively. We also took an attempt to have an influence of mutants in combination with of all single mutant complexes, which led to the formation of same strong hydrogen bonding (L452R; Glu35 at 2.2Å and E484Q; Glu75 at 2.1 Å) with a loss of interaction with T478K. In addition, the Glu35 (hACE2) in L452R consistently sharing its projection with strong hydrogen bonding Arg452, Gln493 and Ser494, while Ser494 equally sharing with Asp38. Ser494, Gly496, Tyr505 and Gln498.

**Figure 3:**
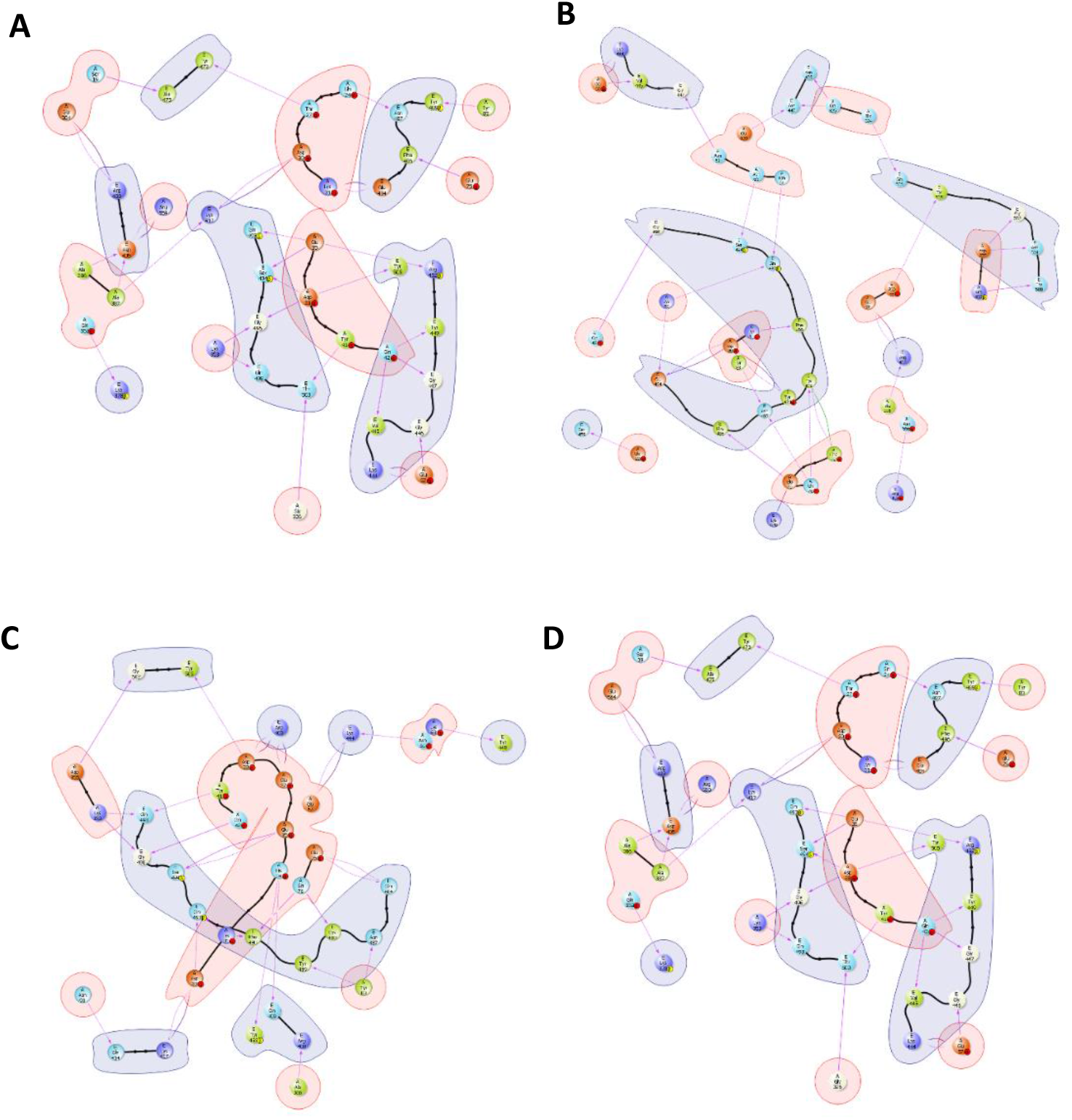
Molecular overlay representing intermolecular contacts in between hACE2 (in orange) and S protein (in greyish) of (A) L452R, (B) T478K, (C) E484Q and (D) L452R + T478K + E484Q mutant complexes.

The mutant complex T478K in the B.1.617.2 lineage, a salt bridge observed in between Glu75 and Lys478 and strong hydrogen bonding with Phe486. Subsequently, side chains gave contacts with Gln493 with Asn64 and Lys68 and non-mutate residue of E484. The structural interface analysis displayed, many polar contacts with Cα backbone possessing stronger affinity thus leads to the evasion of immune escape which supports our results with a reported agreement (Planas et al., 2021). To understand the effect on the interaction patterns of mutant complexes are evaluated and compared with that of wild type. From the interface analysis, E484 residue present on the flexible loop on the RBD projected with Lys31 (hACE2) for the formation of salt bridge. Studies reported that the K31 and K353 are the virus binding hotspots situated on the hACE2 surface (Wan et al., 2022). From the analysis of all complexes, Lys31 and Glu35 projected towards Gln493, while Asp38 and Lys353 gave formation of strong hydrogen bond with Gln498. This supports us the gain in binding affinities of SARS-CoV-2 over SARS-CoV due to the presence of salt bridge between Lys31 (hACE2) and Glu484 (S) (Wrobel et al., 2020). Recently, a study shown mutant E484Q shown increase in affinity over wild type (Yi et al., 2020). To support with this assertion, crystallographic structure proven to have in both closed and open conformations, where open confirmation is only responsible for the receptor binding. Moreover, Glu484 exhibited promising hydrogen bonding contacts with Phe490, while weakens after its mutation. Due to its weakening in contacts, probably it could favour open conformation resulting stronger hACE2 binding and immune escape. The orientation in structural changes, in the RBD due to E484Q mutation might impact on the protein stability binding with hACE2 with help of few experimental reported studies (Cherian et al., 2021; Greaney et al., 2021). To gain deeper understanding, how RBD region in spike protein influence the binding affinity upon the interaction with hACE2, was estimated using extracted frames form all complexes based on the MM-GBSA approach. The binding free energy of the combination of all three mutant simulations were −272.82 kcal/mol. In the single mutant complex involving L452R, T478K and E484Q mutant, the ΔGbind were −189.87 kcal/mol, −226.97 kcal/mol and −187.17 kcal/mol respectively. The comparative analysis revealed that the all mutants exhibited more favourable binding affinity over the wild type of - 59.93 kcal/mol.

In our research highlights, our study relied on the mechanistic structural dynamics approach between hACE2 and S-RBD motif interface of the rapidly spreading B.1.617 lineage using molecular dynamics simulations approach. We attempted on both individual and in combination of mutant(s) to identify and estimate the differences in contacts and alteration in structural conformations of the spike protein over the wild type. Upon the emergence of the variants of concerns from the B.1.617 lineage, amid crisis arose with lot of causalities. The influence of these new variants based on the transmissibility and infectivity among hosts’ cells studied in this research. The results presented in this article performed based on the molecular dynamics and predicted binding affinity in between hACE2 and spike protein. From the results analysis, the mutation E484Q favours open confirmation of RBD that aids better binding and greater immune escape over L452R.

## AUTHOR CONTRIBUTIONS

Both S.A. and J.P. written, performed the computational research and analysed the data; S.A. conceived the research idea and edited the manuscript.

